# Transferrin participates in the pathogenesis of endometriosis by influencing the proliferation, migration and apoptosis of endometrial cells

**DOI:** 10.64898/2026.03.13.711522

**Authors:** Fang Jie, Chaochao Xu, Yunqin Ni, Na Ding, Xinyue Zhang, Haitao Pan

**Author notes:** **Corresponding Author** Haitao Pan: Shaoxing Maternity and Child Health Care Hospital, No. 222 Fenglin East Road, Shaoxing, 312000, Zhejiang, China.

## Abstract

Ferroptosis is linked to various diseases, but the role of transferrin (TF) in endometriosis (EM) remains unclear. Expression levels of ferroptosis-related proteins, including transferrin (TF), transferrin receptor (TFRC), and glutathione peroxidase 4 (GPX4), were analyzed by western blotting. Compared to normal endometrial stromal cells, eutopic and ectopic endometrial stromal cells from EM patients exhibited significantly enhanced proliferative and migratory abilities, accompanied by a marked reduction in glutathione (GSH) levels in both eutopic and ectopic tissues. TF and TFRC expression was upregulated in ectopic endometrium relative to normal controls, while GPX4 expression was downregulated. To evaluate the functional role of TF, siRNA-mediated knockdown was performed in endometrial stromal cells, with knockdown efficiency confirmed by western blotting. Functional assays demonstrated that TF knockdown not only suppressed cell proliferation (CCK-8 and clonogenic assays) and migration (wound healing assay) but also significantly increased apoptosis rate (flow cytometry with Annexin V-FITC/PI staining).These findings implicate TF in the pathogenesis and progression of endometriosis, likely through modulating endometrial stromal cell proliferation, migration, and apoptosis.

## Introduction

Endometriosis (EM) is a condition in which functional endometrial tissue grows and undergoes cyclic bleeding outside the uterine cavity, leading to the formation of nodules, masses, and other types of lesions. It primarily presents as progressively worsening pelvic adhesions, pain, infertility, and other symptoms, making it a common yet challenging gynecological disorder, with an incidence as high as 10% among women of reproductive age[1]. Although histologically benign, EM lesions are widely distributed and morphologically diverse, displaying malignant tumor-like characteristics such as implantation, invasion, recurrence, and metastasis[2], all of which severely affect patients’ health and quality of life. At present, the pathogenesis of EM remains incompletely understood. Studies suggest that abnormal regulation of cellular functions and modes of cell death may be involved in its development and progression.

Hemorrhage in ectopic endometriotic lesions and hemolysis resulting from retrograde menstruation can lead to the release and accumulation of substantial quantities of free iron. Subsequently, intracellular free iron can catalyze the generation of lipid reactive oxygen species (ROS) via the Fenton reaction cascade, resulting in cellular injury. This process is widely recognized as ferroptosis, a novel form of regulated cell death distinct from accidental cell death. Despite many unresolved questions in ferroptosis research, several studies have demonstrated its crucial role in the pathogenesis of various diseases, including endometriosis (EM) [3-6]. Previous investigations have elucidated the complex role of ferroptosis in EM, underscoring its significance in disease development and progression [7-8].On one hand, endometriotic lesions exhibit resistance to ferroptosis, hindering the clearance of ectopic endometrium and facilitating its proliferation and migration [8-9]. On the other hand, ferroptotic cell death can also trigger the release of inflammatory cytokines and activate downstream regulatory pathways, which in turn promote proliferation and angiogenesis in adjacent tissues [10]. The key mechanisms underlying ferroptosis are closely associated with disturbances in iron metabolism, lipid metabolism, and glutathione metabolism [11–13].

Transferrin (Tf), which naturally binds Fe^3^□, serves as the primary iron carrier in the blood and plays a crucial role in iron metabolism and ferroptosis [14]. It is widely distributed across various tissues and organs, where it is closely associated with cell growth and differentiation, and implicated in the pathogenesis of numerous diseases. Previous studies have indicated that abnormal Tf expression is closely linked to the development of several malignancies, including ovarian, breast, liver, and prostate cancers [15–18]. However, the role of Tf in the pathogenesis of endometriosis remains unclear. This study aims to examine the expression of Tf in endometriosis and to explore the impact of its abnormal expression on the biological behavior of endometrial stromal cells.

## Materials and Methods

### Patients and samples

Forty-three participants were enrolled in Shaoxing Maternity and Child Health Care Hospital from January 2022 to September 2024. Control eutopic endometrium (Ctrl) represented samples from non-endometriosis patients. Eutopic endometrium (EuE), and ectopic ovarian lesions (EcO) were collected from revised ASRM Stage II-IV endometriosis patients. All the patients selected for the study had no history of immune disorders, acute inflammatory states, or estrogen-dependent diseases, and had abstained from any hormonal medications for three months prior to enrollment.

### Primary cells isolation and culture

Primary endometrial stromal cells were isolated and cultured as previously described [19]. In brief, tissue specimens were washed with phosphate-buffered saline (PBS), minced into small fragments, and digested with 1 mg/mL collagenase type IV (Sangon Biotech, China) for 20–40 minutes at 37°C on a shaker. The resulting homogenate was filtered through a 40-μm cell strainer (Beyotime Biotechnology, China), and the filtrate was centrifuged for 10 minutes to pellet the cells. The harvested primary cells were resuspended in complete DMEM/F12 medium (Grand Island Biological Company, USA) supplemented with 10% fetal bovine serum (Sangon Biotech, China) and maintained at 37°C in a humidified incubator with 5% CO□. Cells at passage 3 were used for subsequent experiments.

### Western blotting

Total protein from tissues and cells was extracted using RIPA lysis buffer supplemented with phosphatase inhibitors. Cytoplasmic and nuclear proteins were fractionated using a commercial extraction kit (Beyotime Biotechnology, China) according to the manufacturer’s instructions. Protein concentrations were determined by the BCA method. Equal amounts of protein samples were separated by SDS-polyacrylamide gel electrophoresis and subsequently transferred to PVDF membranes (Beyotime Biotechnology, China). After blocking with non-fat milk, the membranes were incubated with primary antibodies against the target proteins overnight at 4°C on a shaker, followed by incubation with an HRP-conjugated secondary antibody. Protein bands were finally visualized using an electrochemiluminescence (ECL) detection system.

### Glutathione (GSH) Assay

Tissue homogenates were prepared by adding protein remover at a ratio of 1:9 (weight g: volume mL) as per the instructions of the GSH assay kit (Beyotime Biotechnology, China). After centrifugation at 10,000 rpm for 15 min, the supernatants were collected for subsequent assays. For the analysis of cultured cells, approximately 1×10□ cells were resuspended and subjected to lysis on ice for 10 min using 100 μL of protein scavenger, followed by centrifugation under identical parameters. The collected supernatant was incubated with the glutathione detection working solution for 5 min. Subsequently, 20 μL of substrate was added and the mixture was vortexed thoroughly. Following a 20-min incubation period, the absorbance at 412 nm was recorded.

### Cell transfection

To knockdown TF, endometrial stromal cells were infected with a specific lentivirus (RiboBio, China), following the supplier’s instructions. Cells were also transfected with the CON520-AURKA plasmid (RiboBio, China) and cultured in serum-containing complete medium at 37°C. After collection and three PBS washes, total RNA was extracted using the TIANGEN DP431 kit. Knockdown efficiency was confirmed by measuring TF mRNA levels using the one-step qRT-PCR method (TIANGEN FP303 kit). Finally, real-time cellular analysis (RTCA) was employed to assess the functional impact of TF knockdown on cell proliferation and migration.

### Cell proliferation assay

Cell proliferation was determined by the CCK-8 assay. Briefly, 96-well plates were seeded with 3,000 cells per well in 100 μL of suspension. After the designated treatments, the cultures were supplemented with 10 μL of CCK-8 reagent (Zeta Life Sciences Inc, UK) at 0, 24, 48, and 72 h. After 4 hours of incubation at 37°C in a 5% CO□ atmosphere, the absorbance at 450 nm was measured with a microplate reader.

### Wound healing assay

A standard wound healing assay was performed. Briefly, cells were cultured in 6-well plates until 80–90% confluent. A linear wound was then introduced across the cell monolayer using a 200 μL pipette tip. After removing cell debris with PBS washes, the culture medium was replaced with serum-free medium. To assess cell migration, the wound areas were photographed at 0, 24, and 48 hours, and the degree of wound closure was quantified by measuring the residual area at these time points.

### Flow cytometric analysis

For apoptosis analysis, endometrial stromal cells (ESCs) were stained with an Annexin V-FITC/PI kit (Yeasen, China) in accordance with the manufacturer’s protocol. The stained cells were then subjected to flow cytometric analysis on an LSR II instrument to determine the proportion of early (Annexin V□/PI□) and late (Annexin V□/PI□) apoptotic cells. Data were processed using FlowJo software (version 10; FlowJo, LLC).

### Statistical analysis

Statistical analyses were conducted with GraphPad Prism, version 6.02. Quantitative data from three independent replicates are expressed as mean ± SD. Differences between groups were assessed by Student’s t-test or one-way ANOVA with appropriate post-hoc tests, and statistical significance was assigned to results with P < 0.05.

## Results

### Cell Proliferative and Migratory Capacities of EM are enhanced

We performed primary cell culture of endometrium cells in the EuE group, the EcO group and the Ctrl group. CCK-8 results showed that the proliferative capacity enhanced in the EuE group and the EcO group compared to the Ctrl group (Figure1.a). Migration is a fundamental property of cells that occurs during many physiological and pathological processes including repair of damaged tissue after injury and the spread of cancer [20]. We found that the time required for wound closure of endometrium cells in the EcO group was significantly shorter than the time required for the Ctrl group (Figure1.b). The present results obviously demonstrated that the proliferative and Migratory capacities of endometrium cells are closely related to the occurrence of EM.

**Figure 1.**
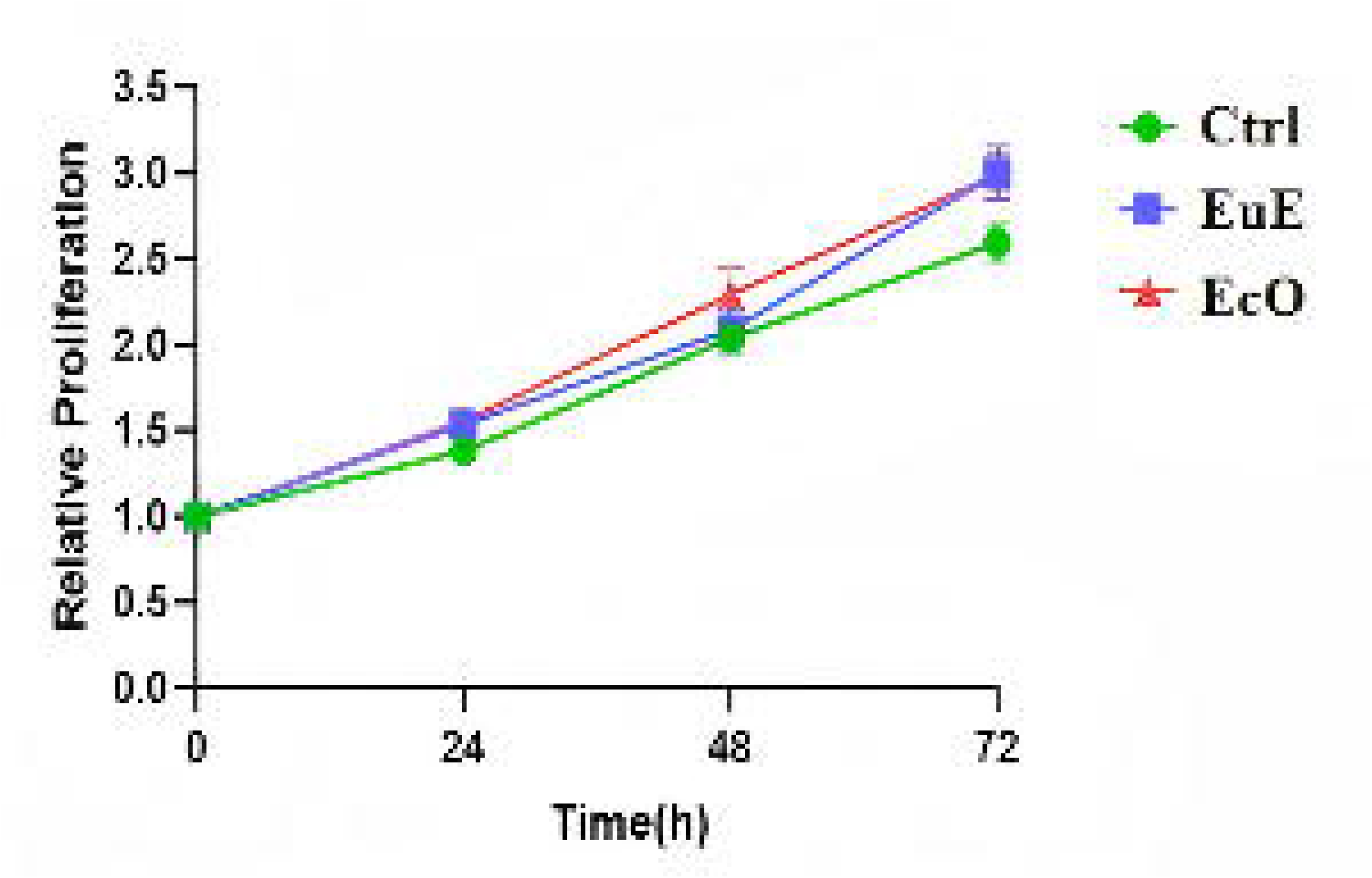
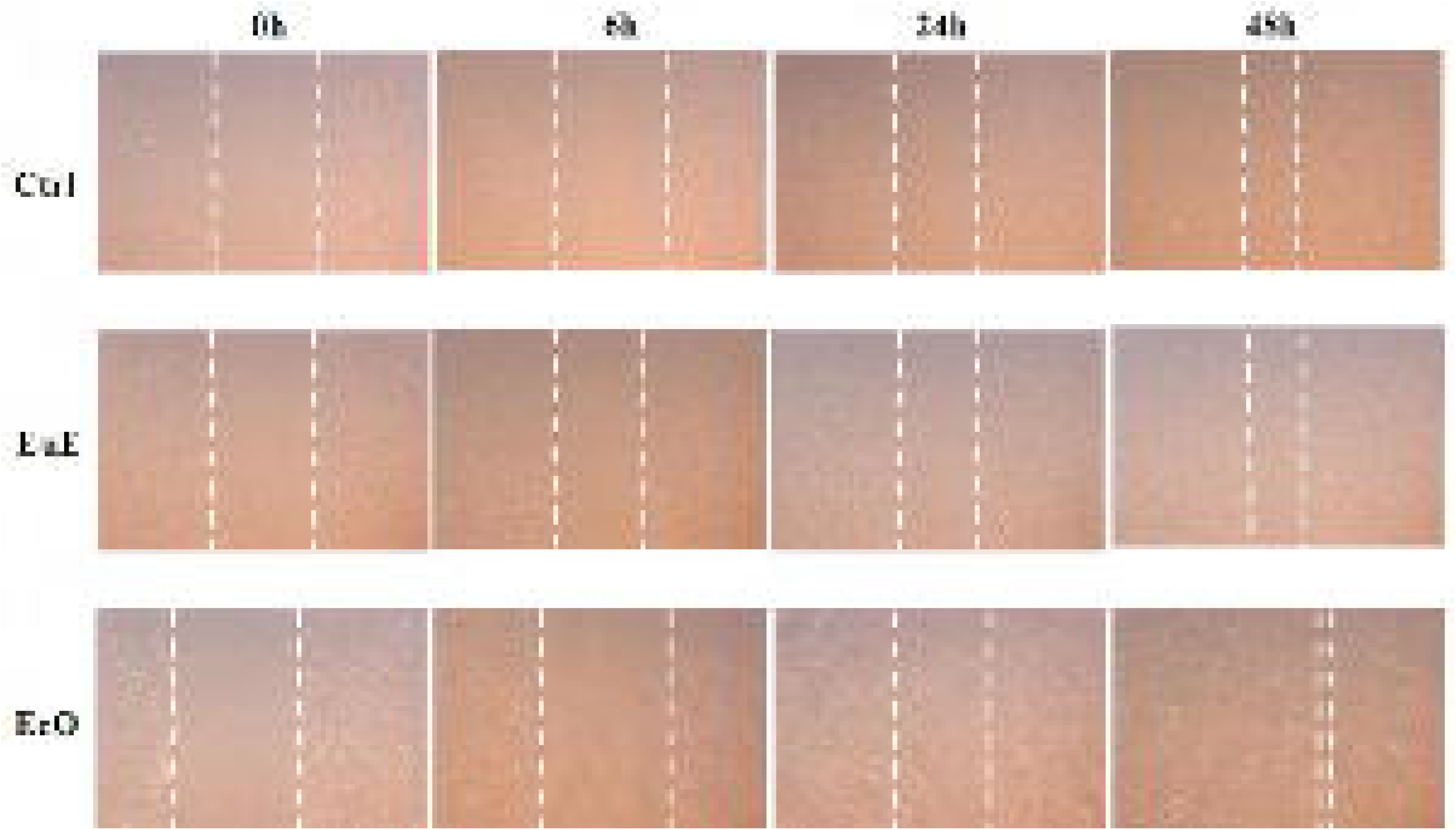
Endometriotic cells exhibit enhanced proliferative and migratory capacities. (a) Cell proliferation was assessed by CCK-8 assay. The eutopic (EuE) and ectopic (EcO) endometrial stromal cells from EM patients showed significantly enhanced proliferation compared to control (Ctrl) endometrial stromal cells after 72 hours of culture. (b) Cell migration was evaluated by wound healing assay. Confluent monolayers of human endometrial stromal cells were wounded with a 200-μL pipette tip and photographed at the indicated time points (0, 6, 24, and 48 hours). The EcO group demonstrated a significantly faster wound closure rate compared to the Ctrl group.

### GSH content is inhibited in EM

GSH is a linear tripeptide of l-glutamine, l-cysteine, and glycine, and is one of the most abundant and significant scavengers of ROS in eukaryotic cells [21]. A feature of ferroptosis is the reduction of antioxidant activity (e.g., intracellular GSH depletion) [22]. As shown in figure 2, the GSH level in the EuE group was much lower, by 97.18% (p<0.001), than that of the Ctrl group. And the GSH level in the EcO group was lower, by 71.13%(p<0.001), than that of the Ctrl group. The present results demonstrated that, GSH is inhibited in EM.

**Figure 2.**
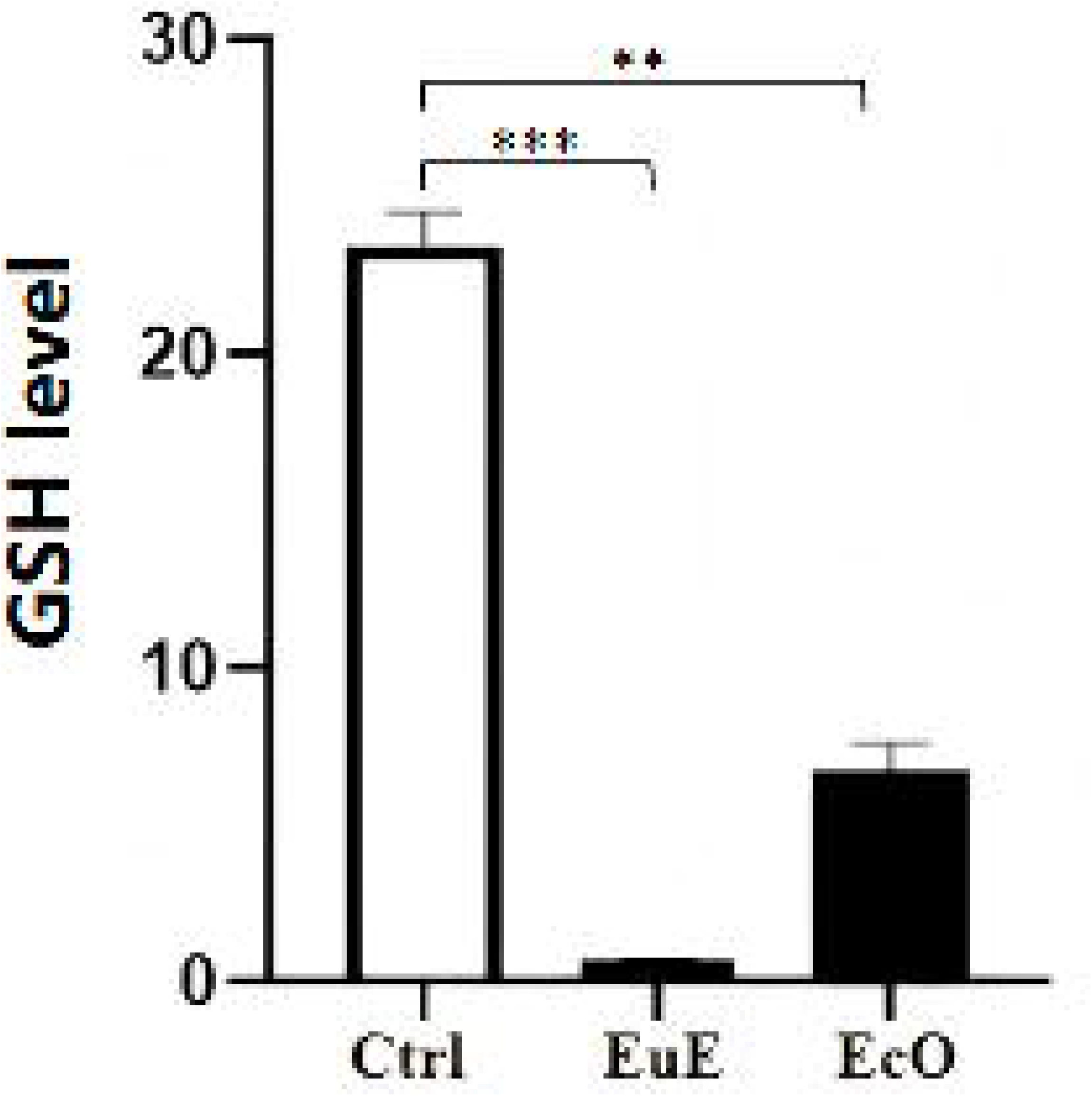
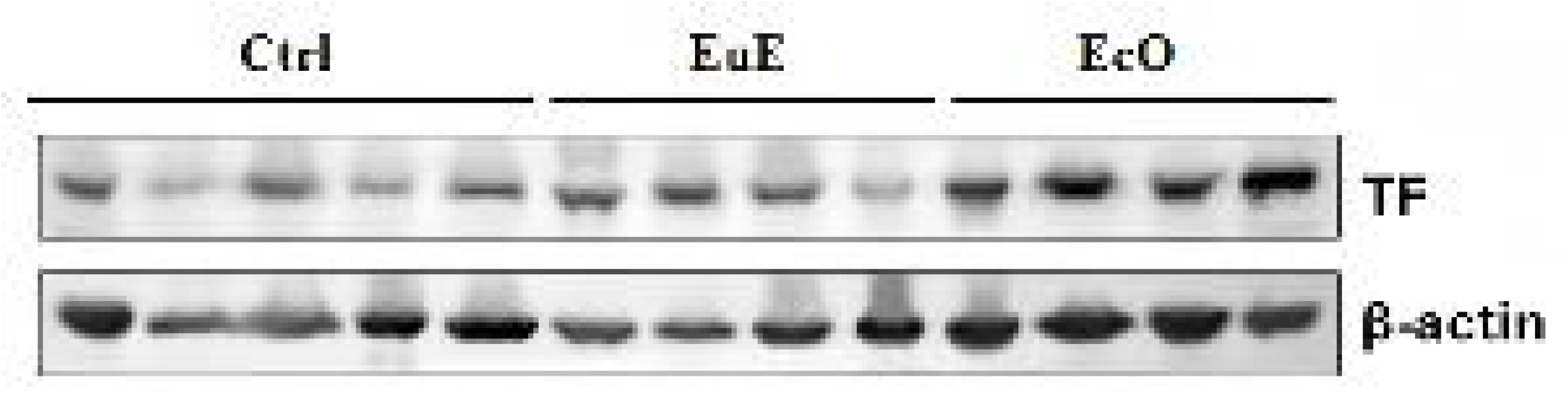
Glutathione (GSH) levels are significantly reduced in endometriosis (EM). The GSH content was substantially decreased in both eutopic (EuE) and ectopic (EcO) endometrial tissues from EM patients compared to the control (Ctrl) group. Data are expressed as the mean ± SEM. *p < 0.05, **p < 0.01, ***p < 0.001; ns, not significant.

### Ferroptosis is Increased in EM

It is well known that GPX4 is the critical repressor of ferroptosis [23]. And TF/TFRC is an iron carrier protein that induces ferroptosis [24]. Western blot were used to detect the expression of TF, TFRC,and GPX4 in the three gtoups from protein levels. As shown in figure 3, TF was significantly upregulated in the EcO group compared to the Ctrl group (p<0.05). TFRC was significantly upregulated in the EuE group compared to the Ctrl group (p<0.05). GPX4 protein expression is downregulated in the EcO group compared to the Ctrl group (p<0.05). To further validate the role of TF in EM, we sought to down-regulate the mRNA expression of TF in endometrium cells using siRNA techniques for the subsequent experiments.

**Figure 3.**
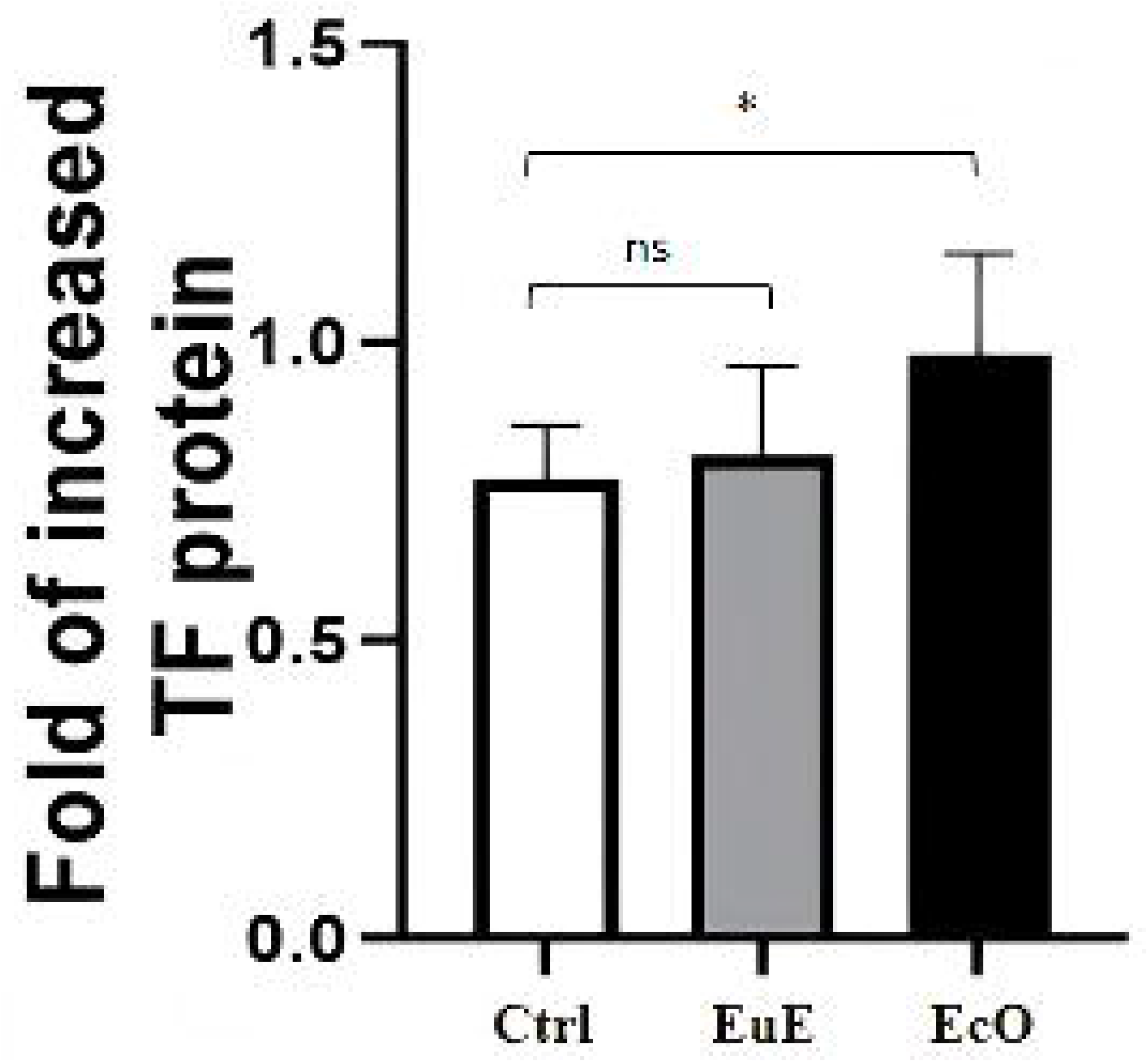
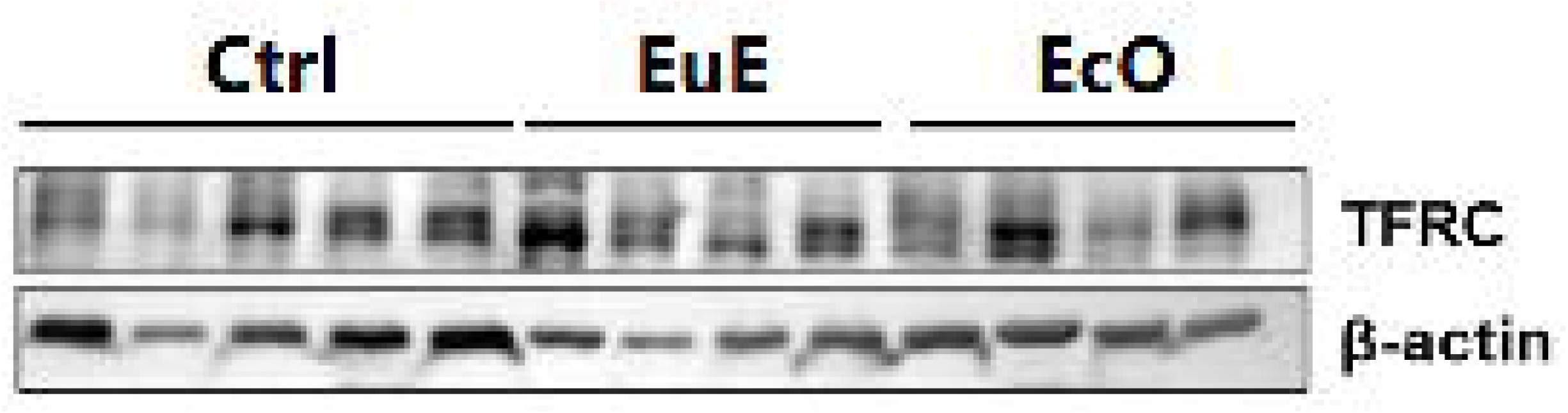
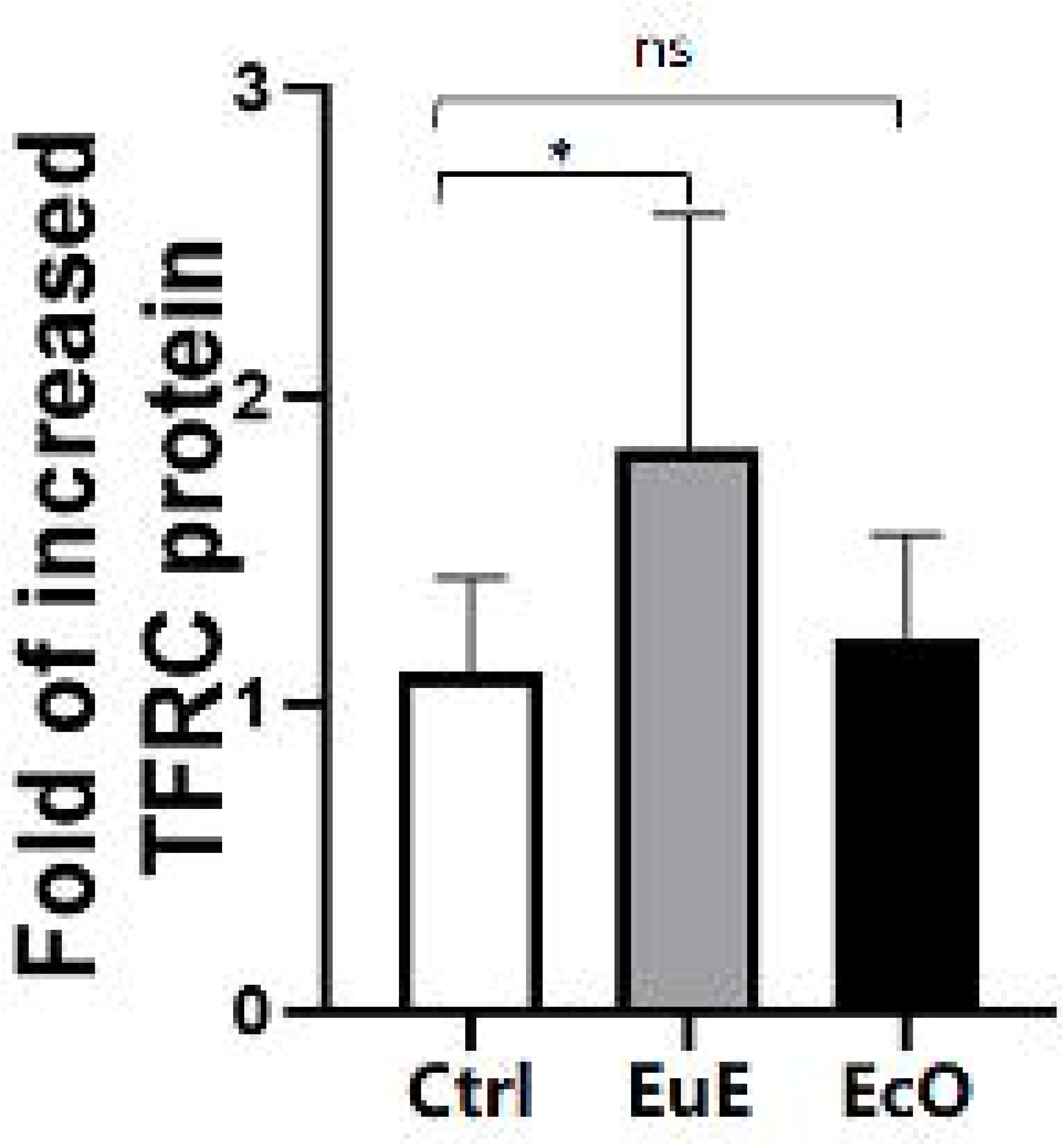
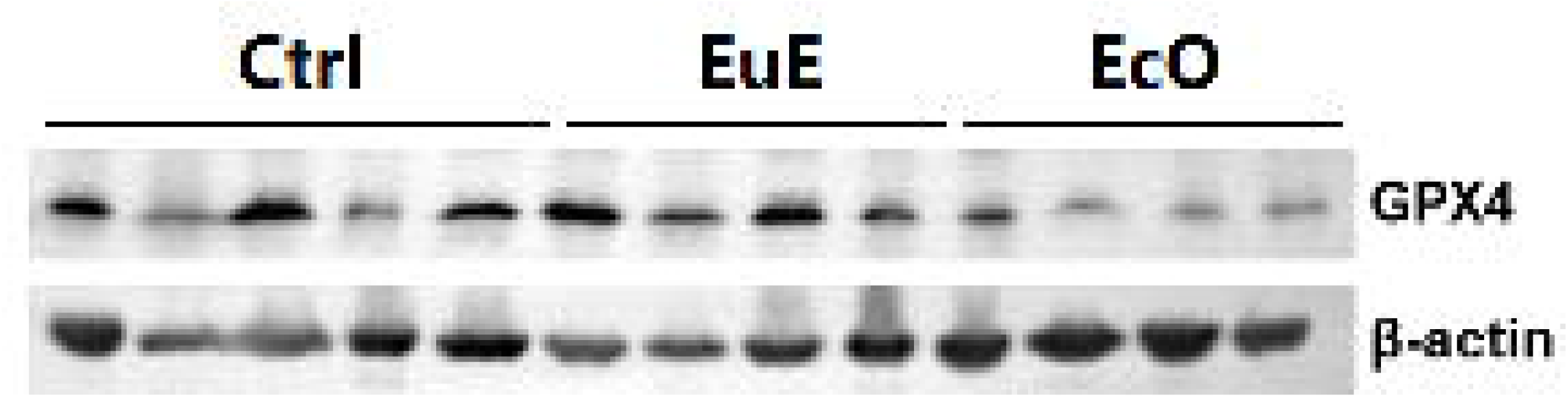
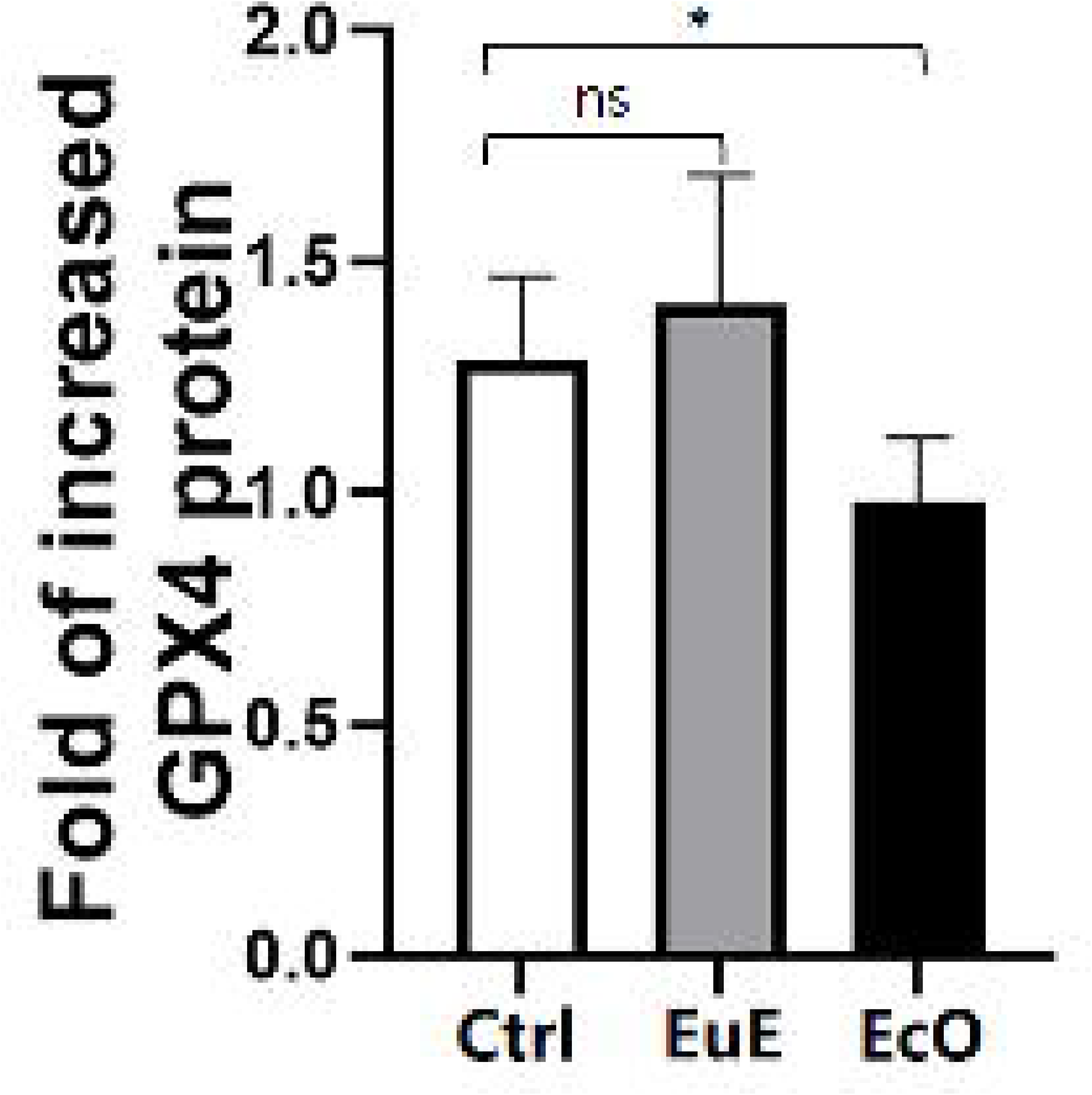
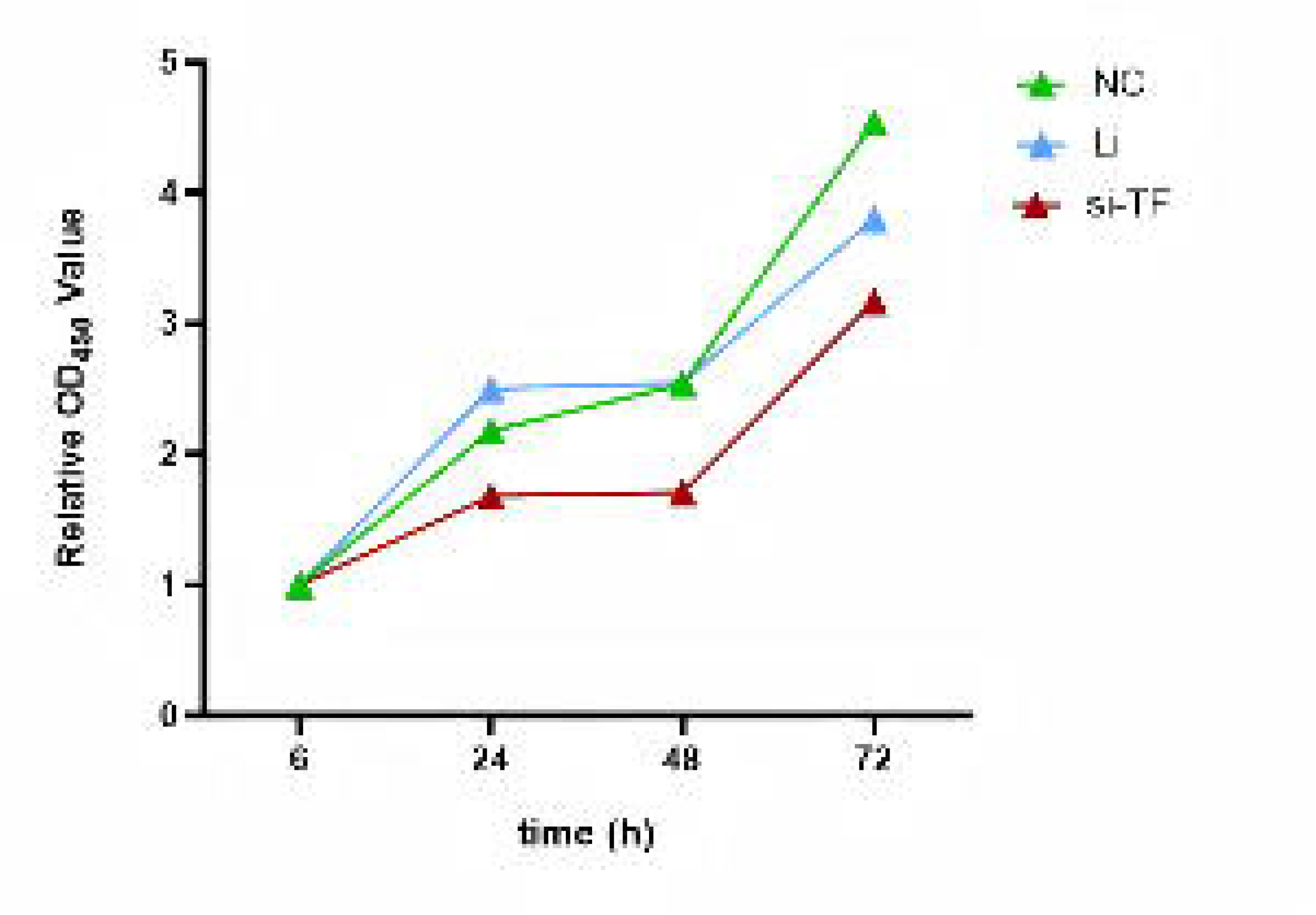
Expression of ferroptosis-related proteins in endometriotic lesions. (a, c, e) Representative western blot images of TF, TFRC, GPX4 and β-actin (loading control) in control (Ctrl), eutopic (EuE), and ectopic (EcO) endometrial tissues. (b, d, f) Quantitative analysis of (b) TF, (d) TFRC, and (f) GPX4 protein levels normalized to β-actin. Quantitative data show that TF expression was significantly higher in the EcO group compared to the Ctrl group, while TFRC was significantly upregulated in the EuE group. GPX4 protein expression was downregulated in the EcO group. Data are expressed as the mean ± SEM. *p < 0.05, **p < 0.01, ***p < 0.001; ns, not significant.

### Knockdown of TF inhibits cell proliferation, migration and apoptosis of endometrium cells after transfection

To investigate the role of TF in endometrium cells, the TF-specific si-TF was designed and transfected into endometrium cells to further determine its effect on the cell growth of endometrium cell in vitro. CCK-8 assay results revealed that the TF knockdown obviously suppressed the proliferation rate of endometrium cells (Figure 4.a). Wound healing assay was used to assess the migration ability of the cells, and the results revealed that, compared with the control conditions, knockdown of TF significantly decreased the wound healing rate and number of migrated cells (Figure 4.b). Flow cytometry was carried out for cell apoptosis examination, the results revealed that, knockdown of TF significantly increased apoptosis rate in endometrium cells (Figure 4.c).

**Figure 4.**
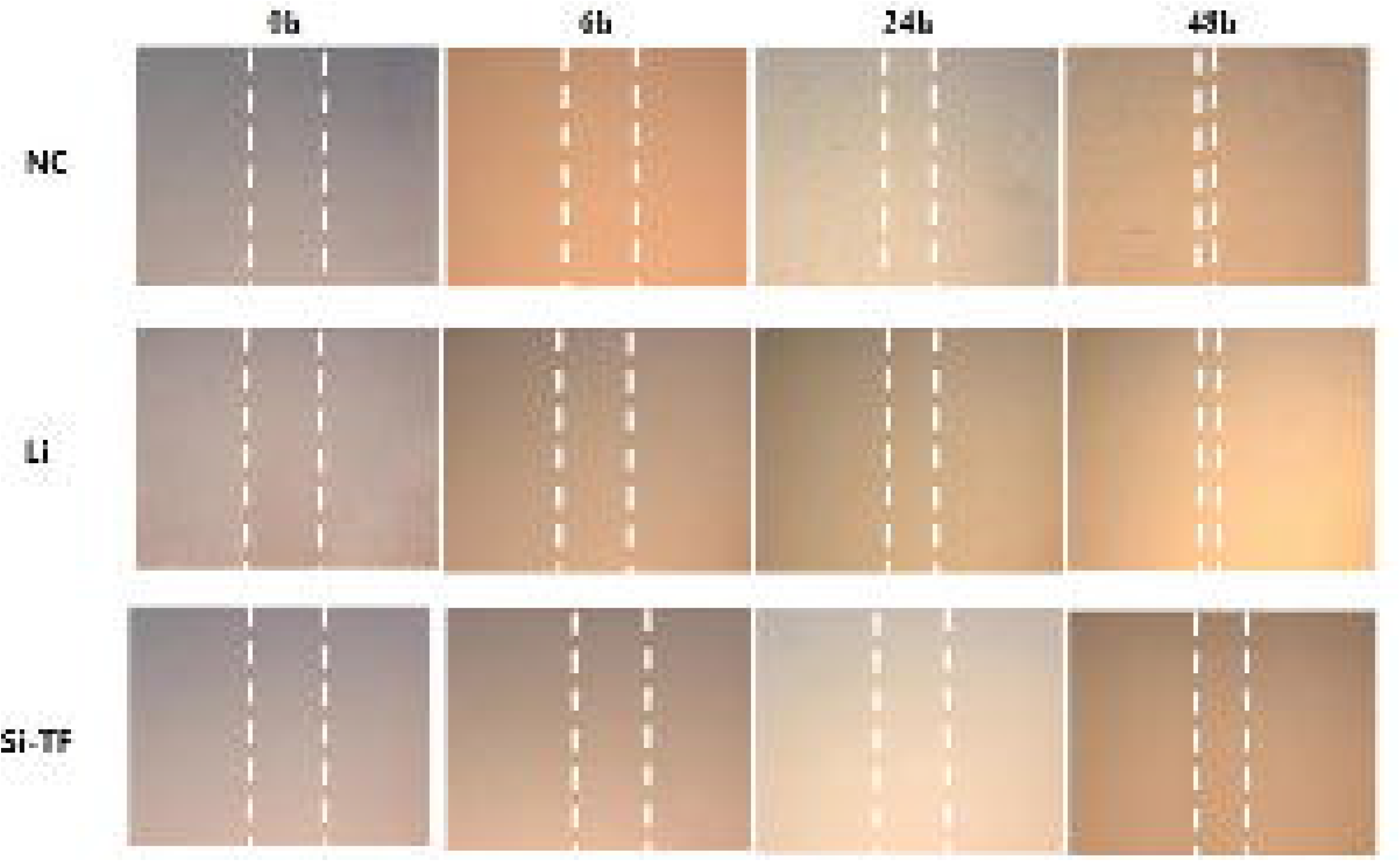
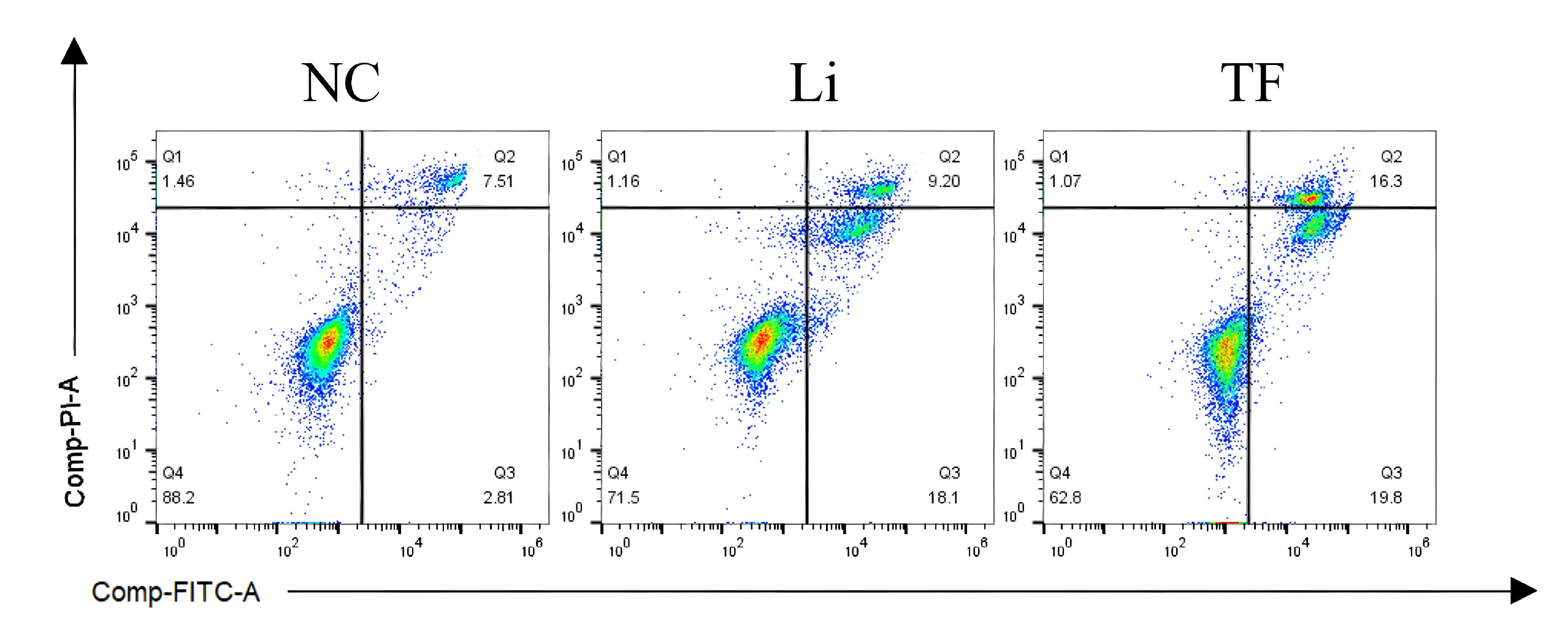
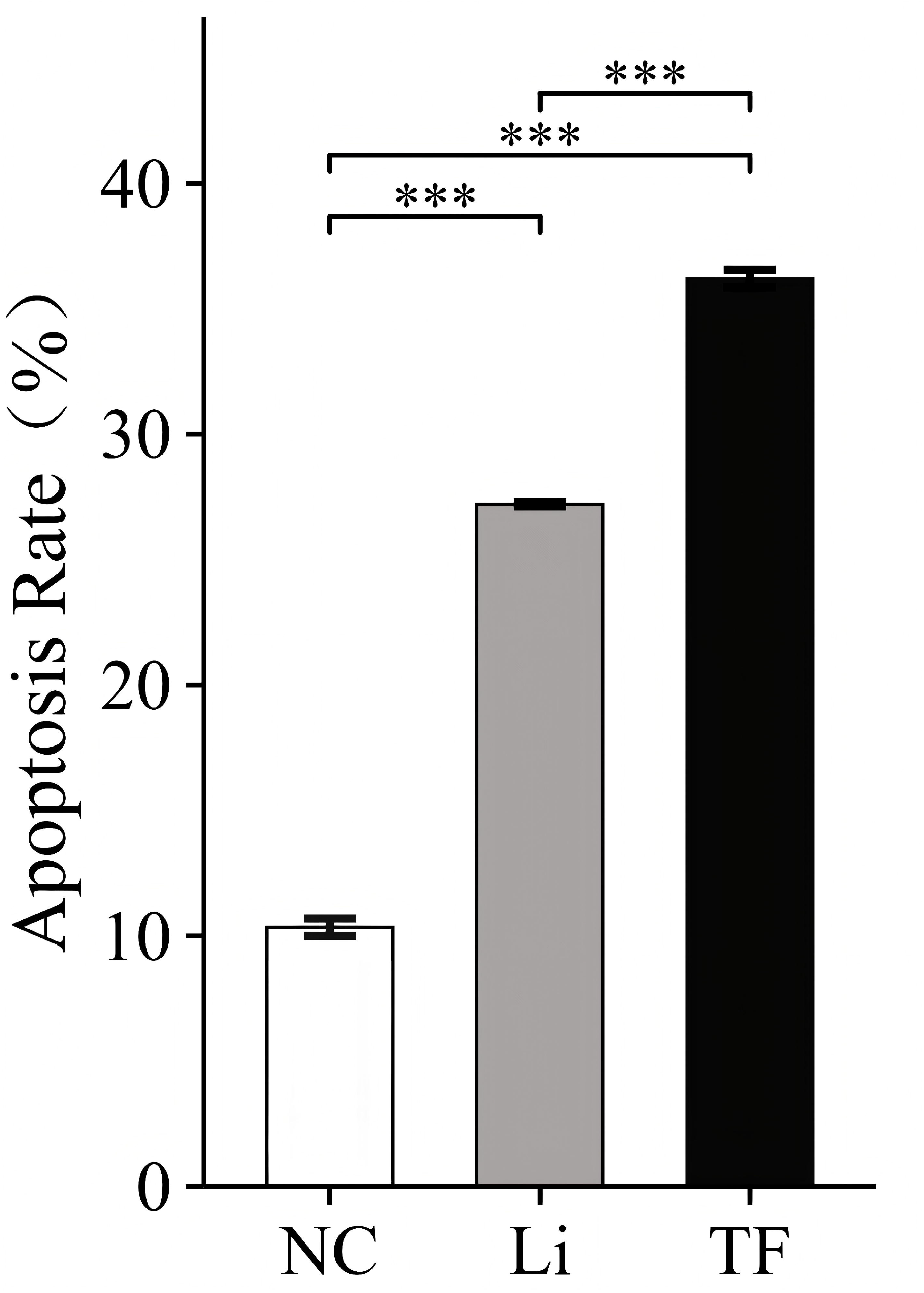
Knockdown of TF inhibits endometrial stromal cells proliferation and migration, increases apoptosis in vitro. (a) The proliferative capacity of endometrial stromal cells was assessed by CCK-8 assay following TF knockdown. Knockdown of TF significantly suppressed cell proliferation compared to the control groups. (b) Cell migration was evaluated by wound healing assay. Confluent monolayers of the indicated endometrial stromal cells groups were scratched and monitored over 96 hours. TF knockdown markedly delayed wound closure and reduced the number of migrated cells relative to controls. (c,d) Apoptosis was assessed by flow cytometry. TF knockdown significantly increased the apoptosis rate of endometrial stromal cells compared to control groups. Data are expressed as the mean ± SEM. *p < 0.05, **p < 0.01, ***p < 0.001; ns, not significant.

## Discussion

The precise pathogenesis of endometriosis, though not yet fully elucidated, is recognized to involve aberrant sex-steroid signaling and inflammatory processes. The retrograde flux of endometrial tissue into the pelvic cavity during menstruation represents a well-established mechanism for the initiation of pelvic lesions, while lymphatic or vascular metastasis has been suggested as a potential pathway for the development of extrapelvic lesions [24-26]. Previous research by Chinese scholar Jinghe Lang and colleagues further identified fundamental abnormalities in the eutopic endometrium of women with endometriosis, which exhibits enhanced capacities for proliferation, implantation, and angiogenesis, along with a heightened ability to survive in ectopic microenvironments [27]. In line with these observations, our study demonstrated that eutopic endometrial stromal cells from EM patients display significantly increased proliferative and migratory activities compared to normal endometrial stromal cells from non-EM individuals, with these aberrant behavioral characteristics being even more pronounced in ectopic lesions. These findings collectively indicate that functional alterations in endometrial stromal cells are closely associated with the pathogenesis of endometriosis.

Hemorrhage in ectopic endometriotic lesions and hemolysis resulting from retrograde menstruation lead to the release and accumulation of substantial amounts of insoluble Fe^3^□. Subsequent biochemical reactions, notably the Fenton reaction, drive the generation of reactive oxygen species (ROS), which induce lipid peroxidation. Consistent with this mechanism, studies have reported that patients with endometriosis exhibit decreased antioxidant system activity alongside significantly elevated levels of ROS and lipid peroxidation markers in both serum and peritoneal fluid compared to healthy women [28]. The persistent production and accumulation of ROS inflict considerable damage on cellular structures and functions, promoting somatic mutations, tumorigenic transformation, and proliferative responses. Moreover, ROS are intimately involved in metabolic and proliferative signaling, with dysregulated ROS pathways being implicated in cancer progression and chronic inflammatory diseases [29–31]. To maintain redox homeostasis and mitigate oxidative damage, the body employs a sophisticated antioxidant system. Among its components, glutathione (GSH) serves a critical role in antioxidative defense, protecting cells from oxidative stress in the context of endometriosis [32]. This system is critically coordinated by glutathione peroxidase 4 (GPX4) [23], which utilizes GSH to specifically reduce lipid hydroperoxides, thereby preventing the iron-dependent peroxidative chain reactions that drive ferroptosis. In our study, we observed a significant reduction in GSH content in both eutopic and ectopic endometrial tissues from EM patients relative to normal endometrium. Consistently, GPX4 protein expression was also significantly downregulated in ectopic lesions. The concurrent reduction in both GSH and GPX4 suggests a weakened anti-ferroptotic capacity, potentially increasing the susceptibility of endometriotic cells to iron-dependent oxidative damage and ferroptosis.

The primary mechanism for cellular iron uptake is mediated by transferrin (TF), a liver-synthesized glycoprotein that functions as a natural chelator with two high-affinity binding sites for ferric iron (Fe^3^□) [33–34]. In the context of endometriosis, the release and accumulation of substantial Fe^3^□ from ectopic lesions allow TF to bind these ions. The resulting transferrin-bound iron is internalized via binding to transferrin receptor (TFRC) and clathrin-mediated endocytosis. During this process, Fe^3^□ is released, reduced to Fe^2^□, and transported into the cytoplasm through divalent metal transporter 1 (DMT1), contributing to the intracellular labile iron pool [35–36]. Subsequently, cytosolic Fe^2^□ catalyzes the Fenton reaction, thereby driving lipid peroxidation of cellular membranes and generating lipid-derived ROS. This oxidative stress is implicated in diverse pathological processes, including chronic inflammation and cancer [37–38]; furthermore, ROS play an established role in promoting tumor angiogenesis, migration, invasion, and metastasis [39–41].In this study, our observation of elevated TF and TFRC expression in ectopic endometrium from EM patients, relative to normal endometrium, suggests a model in which increased TF-mediated iron import enhances intracellular labile iron availability, thereby stimulating ROS production and influencing cellular behavior. To further investigate the functional role of TF, we performed TF knockdown and observed a corresponding decrease in proliferative and migratory capacities, along with a significant increase in the apoptosis rate. These results imply that aberrant TF expression may contribute to the pathogenesis and progression of endometriosis by modulating key cellular behaviors such as proliferation, migration and apoptosis.

## Conclusion

In summary, this study demonstrates that TF is dysregulated and highly expressed in endometriosis. Knockdown of TF inhibits disease progression, likely by modulating ferroptosis and apoptosis pathways, which provides new evidence for TF as a potential therapeutic target. However, further validation in animal models and clinical studies is required.

## Ethics approval and consent to participate

This study was approved by the Medical Ethics Committee of Shaoxing Maternity and Child Health Care Hospital (Approval No. 2022-001).

## Consent for publication

Written informed consent was obtained from all individual participants involved in the study. Participants consented to the publication of anonymized data derived from their samples and clinical information. This study was conducted in accordance with the ethical standards of the institutional research committee.

## Availability of data and materials

The datasets generated and/or analysed during the current study are not publicly available due to the sensitive nature of the clinical data involved and to protect the privacy of the participants, but are available from the corresponding author (Haitao Pan) on reasonable request.

## Authors’ contributions

FJ designed the study, mainly completed the experiment, and wrote the manuscript under the direction of HTP. CCX participated in study design and advised the paper. YQN, ND and XYZ performed the experiments and analyzed data. HTP contributed to experimental design, interpretation of results, and article revision.

## Acknowledgments

We sincerely thank all the personnel involved in this study for their valuable contributions.

## References

1. As-Sanie, S., et al., Endometriosis: A Review. JAMA, 2025. 334(1): p. 64–78.DOI: 10.1001/jama.2025.2975.

2. Asghari, S., et al., Endometriosis: Perspective, lights, and shadows of etiology. Biomed Pharmacother, 2018. 106: p. 163–174.DOI: 10.1016/j.biopha.2018.06.109.

3. Jiang, X., B.R. Stockwell, and M. Conrad, Ferroptosis: mechanisms, biology and role in disease. Nat Rev Mol Cell Biol, 2021. 22(4): p. 266–282.DOI: 10.1038/s41580-020-00324-8.

4. Yan, H.F., et al., Ferroptosis: mechanisms and links with diseases. Signal Transduct Target Ther, 2021. 6(1): p. 49.DOI: 10.1038/s41392-020-00428-9.

5. Bell, H.N., B.R. Stockwell, and W. Zou, Ironing out the role of ferroptosis in immunity. Immunity, 2024. 57(5): p. 941–956.DOI: 10.1016/j.immuni.2024.03.019.

6. Ng, S.W., et al., Endometriosis: The Role of Iron Overload and Ferroptosis. Reprod Sci, 2020. 27(7): p. 1383–1390.DOI: 10.1007/s43032-020-00164-z.

7. Li, B., et al., Ferroptosis resistance mechanisms in endometriosis for diagnostic model establishment. Reprod Biomed Online, 2021. 43(1): p. 127–138.DOI: 10.1016/j.rbmo.2021.04.002.

8. Li, Y., et al., Double-edged roles of ferroptosis in endometriosis and endometriosis-related infertility. Cell Death Discov, 2023. 9(1): p. 306.DOI: 10.1038/s41420-023-01606-8.

9. Wan, Y., et al., Long noncoding RNA ADAMTS9-AS1 represses ferroptosis of endometrial stromal cells by regulating the miR-6516-5p/GPX4 axis in endometriosis. Sci Rep, 2022. 12(1): p. 2618.DOI: 10.1038/s41598-022-04963-z.

10. Li, G., et al., Endometrial stromal cell ferroptosis promotes angiogenesis in endometriosis. Cell Death Discov, 2022. 8(1): p. 29.DOI: 10.1038/s41420-022-00821-z.

11. Yang, Y., et al., Interaction between macrophages and ferroptosis. Cell Death Dis, 2022. 13(4): p. 355.DOI: 10.1038/s41419-022-04775-z.

12. Ou, M., et al., Role and mechanism of ferroptosis in neurological diseases. Mol Metab, 2022. 61: p. 101502.DOI: 10.1016/j.molmet.2022.101502.

13. Tang, D., et al., Ferroptosis: molecular mechanisms and health implications. Cell Res, 2021. 31(2): p. 107–125.DOI: 10.1038/s41422-020-00441-1.

14. Gao, M., et al., Glutaminolysis and Transferrin Regulate Ferroptosis. Mol Cell, 2015. 59(2): p. 298–308.DOI: 10.1016/j.molcel.2015.06.011.

15. Deshpande, P., et al., Transferrin and octaarginine modified dual-functional liposomes with improved cancer cell targeting and enhanced intracellular delivery for the treatment of ovarian cancer. Drug Deliv, 2018. 25(1): p. 517–532.DOI: 10.1080/10717544.2018.1435747.

16. Li, L., et al., Targeted Delivery of Doxorubicin Using Transferrin-Conjugated Carbon Dots for Cancer Therapy. ACS Appl Bio Mater, 2021. 4(9): p. 7280–7289.DOI: 10.1021/acsabm.1c00811.

17. Huang, S.Z., et al., Targeting TF-AKT/ERK-EGFR Pathway Suppresses the Growth of Hepatocellular Carcinoma. Front Oncol, 2019. 9: p. 150.DOI: 10.3389/fonc.2019.00150.

18. Peng, J., et al., Adenoviral Vector for Enhanced Prostate Cancer Specific Transferrin Conjugated Drug Targeted Therapy. Nano Lett, 2022. 22(10): p. 4168–4175.DOI: 10.1021/acs.nanolett.2c00931.

19. Ryan, I.P., E.D. Schriock, and R.N. Taylor, Isolation, characterization, and comparison of human endometrial and endometriosis cells in vitro. J Clin Endocrinol Metab, 1994. 78(3): p. 642–9.DOI: 10.1210/jcem.78.3.8126136.

20. Merino-Casallo, F., et al., Unravelling cell migration: defining movement from the cell surface. Cell Adh Migr, 2022. 16(1): p. 25–64.DOI: 10.1080/19336918.2022.2055520.

21. Zitka, O., et al., Redox status expressed as GSH:GSSG ratio as a marker for oxidative stress in paediatric tumour patients. Oncol Lett, 2012. 4(6): p. 1247–1253.DOI: 10.3892/ol.2012.931.

22. Stockwell, B.R., et al., Ferroptosis: A Regulated Cell Death Nexus Linking Metabolism, Redox Biology, and Disease. Cell, 2017. 171(2): p. 273–285.DOI: 10.1016/j.cell.2017.09.021.

23. Chen, X., et al., Cellular degradation systems in ferroptosis. Cell Death Differ, 2021. 28(4): p. 1135–1148 DOI: 10.1038/s41418-020-00728-1.

24. Horne, A.W. and S.A. Missmer, Pathophysiology, diagnosis, and management of endometriosis. BMJ, 2022. 379: p. e070750.DOI: 10.1136/bmj-2022-070750.

25. Saunders, P.T.K. and A.W. Horne, Endometriosis: Etiology, pathobiology, and therapeutic prospects. Cell, 2021. 184(11): p. 2807–2824.DOI: 10.1016/j.cell.2021.04.041.

26. Zondervan, K.T., C.M. Becker, and S.A. Missmer, Endometriosis. N Engl J Med, 2020. 382(13): p. 1244–1256.DOI: 10.1056/NEJMra1810764.

27. Liu, H. and J.H. Lang, Is abnormal eutopic endometrium the cause of endometriosis? The role of eutopic endometrium in pathogenesis of endometriosis. Med Sci Monit, 2011. 17(4): p. RA92–9.DOI: 10.12659/msm.881707.

28. Scutiero, G., et al., Oxidative Stress and Endometriosis: A Systematic Review of the Literature. Oxid Med Cell Longev, 2017. 2017: p. 7265238.DOI: 10.1155/2017/7265238.

29. Chen, K., et al., Mitochondrial mutations and mitoepigenetics: Focus on regulation of oxidative stress-induced responses in breast cancers. Semin Cancer Biol, 2022. 83: p. 556–569.DOI: 10.1016/j.semcancer.2020.09.012.

30. Deng, Y.Q., et al., Compound-composed Chinese medicine of Huachansu triggers apoptosis of gastric cancer cells through increase of reactive oxygen species levels and suppression of proteasome activities. Phytomedicine, 2024. 123: p. 155169.DOI: 10.1016/j.phymed.2023.155169.

31. Shao, W., et al., Ferroportin inhibits the proliferation and migration of fibroblast-like synoviocytes in rheumatoid arthritis via regulating ROS/PI3K/AKT signaling pathway. Eur J Pharmacol, 2025. 987: p. 177205.DOI: 10.1016/j.ejphar.2024.177205.

32. Kobayashi, H., et al., Current Understanding of and Future Directions for Endometriosis-Related Infertility Research with a Focus on Ferroptosis. Diagnostics (Basel), 2023. 13(11).DOI: 10.3390/diagnostics13111926.

33. Lei, G., L. Zhuang, and B. Gan, Targeting ferroptosis as a vulnerability in cancer. Nat Rev Cancer, 2022. 22(7): p. 381–396.DOI: 10.1038/s41568-022-00459-0.

34. Galy, B., et al., Iron regulatory proteins secure mitochondrial iron sufficiency and function. Cell Metab, 2010. 12(2): p. 194–201.DOI: 10.1016/j.cmet.2010.06.007.

35. Muckenthaler, M.U., et al., A Red Carpet for Iron Metabolism. Cell, 2017. 168(3): p. 344–361.DOI: 10.1016/j.cell.2016.12.034.

36. Jabara, H.H., et al., A missense mutation in TFRC, encoding transferrin receptor 1, causes combined immunodeficiency. Nat Genet, 2016. 48(1): p. 74–8.DOI: 10.1038/ng.3465.

37. Climent, M., et al., MicroRNA and ROS Crosstalk in Cardiac and Pulmonary Diseases. Int J Mol Sci, 2020. 21(12).DOI: 10.3390/ijms21124370.

38. Chen, C., et al., Mitochondria and oxidative stress in ovarian endometriosis. Free Radic Biol Med, 2019. 136: p. 22–34.DOI: 10.1016/j.freeradbiomed.2019.03.027.

39. Lopez-Mejia, I.C., et al., Oxidative stress-induced FAK activation contributes to uterine serous carcinoma aggressiveness. Mol Oncol, 2023. 17(1): p. 98–118.DOI: 10.1002/1878-0261.13346.

40. Fukai, T. and M. Ushio-Fukai, Cross-Talk between NADPH Oxidase and Mitochondria: Role in ROS Signaling and Angiogenesis. Cells, 2020. 9(8).DOI: 10.3390/cells9081849.

41. Wang, L., et al., ROS-producing immature neutrophils in giant cell arteritis are linked to vascular pathologies. JCI Insight, 2020. 5(20).DOI: 10.1172/jci.insight.139163.

